# Light modulates glucose and lipid homeostasis via the sympathetic nervous system

**DOI:** 10.1101/2024.10.11.617839

**Authors:** Xiangning Chen, Eugene Lin, Mozhan M. Haghighatian, Logan Wallace Shepard, Samer Hattar, Rejji Kuruvilla, Haiqing Zhao

## Abstract

Light is an important environmental factor for vision, and for diverse physiological and psychological functions. Light can also modulate glucose metabolism. Here, we show that in mice, light is critical for glucose and lipid homeostasis by regulating the sympathetic nervous system, independent of circadian disruption. Light deprivation from birth elicits insulin hypersecretion, glucagon hyposecretion, lower gluconeogenesis, and reduced lipolysis by 6- 8 weeks, in male, but not, female mice. These metabolic defects are consistent with blunted sympathetic activity, and indeed, sympathetic responses to a cold stimulus are significantly attenuated in dark-reared mice. Further, long-term dark rearing leads to body weight gain, insulin resistance, and glucose intolerance. Notably, metabolic dysfunction can be partially alleviated by 5 weeks exposure to a regular light-dark cycle. These studies provide insight into circadian-independent mechanisms by which light directly influences whole-body physiology and inform new approaches for understanding metabolic disorders linked to aberrant environmental light conditions.

**Teaser:** Light exerts direct circadian-independent effects on glucose and lipid metabolism.

## Introduction

Light is one of the most important environmental signals on earth affecting health, survival, and well-being in animals by influencing image-forming vision and a host of non-image functions. Exposure to a regular light-dark cycle has profound effects on circadian photoentrainment, mood, and cognition (*1*), as well as maintaining physiological homeostasis through the regulation of sleep, glucose metabolism, body temperature, and energy expenditure (*2–5*). In mammals, light detection occurs predominantly in the retina and requires the photoreceptors, rods, cones, and the more recently discovered, intrinsically photosensitive retinal ganglion cells (ipRGCs), that express the photopigment, melanopsin (*6*). In particular, ipRGCs integrate extrinsic synaptic input from the canonical rod and cone photoreceptors with their intrinsic melanopsin-based photoreceptive system to convey environmental light signals for diverse non-image functions (*1, 7–9*). While it has been assumed that most of these light-mediated effects require the photoentrainment of the circadian clock located in the suprachiasmatic nucleus (SCN), accumulating evidence suggests that light is also capable of directly affecting animal physiology and behavior through pathways independent of circadian rhythms (*10–12*).

The dynamic control of glucose metabolism in response to environmental challenges is vital for the survival of animals. Epidemiologic studies indicate that aberrant light conditions are one of the highest risk factors for metabolic diseases, including insulin resistance, glucose intolerance, diabetes, and obesity (*3, 5, 13, 14*). Altered glucose regulation has also been observed in both humans and rats with congenital blindness (*15, 16*), highlighting the importance of light detection in the retina in regulating glucose metabolism. Misalignment of light exposure with innate circadian clocks is thought to be the primary pathway by which aberrant light conditions affect glucose and energy homeostasis. While most studies have highlighted circadian misalignment as the major culprit underlying metabolic dysfunction, a recent study has reported that a brief light exposure (2 hr) acutely impairs glucose tolerance by inhibiting thermogenesis in mice (*10*). Overall, how light influences metabolic physiology in a circadian-independent manner is vastly understudied.

The sympathetic nervous system is a major conduit for communication between the brain and peripheral tissues in maintaining whole-body homeostasis in the face of external or internal challenges (*17*). In particular, the sympathetic nervous system works cooperatively with the central nervous system (CNS) to regulate blood glucose homeostasis by controlling pancreatic hormone secretion, glucose production in the liver, as well as its uptake by peripheral tissues, including skeletal muscle and adipose tissues (*18–22*). The sympathetic nervous system also regulates energy balance by promoting brown adipose tissue (BAT)- mediated thermogenesis and triggering lipolysis in white adipose tissue (*23*). Light is known to activate the sympathetic nervous system in humans and rodents, based on electrophysiological recordings (*24–26*). Limited studies have also reported that light can directly modulate peripheral organ/tissue functions, including hair follicle stem cell proliferation and brown adipose tissue-mediated thermogenesis, via the sympathetic nervous system (*10, 11*). Whether light influences glucose and lipid metabolism via the sympathetic nervous system in a manner independent of circadian rhythms remains unclear.

Here we used dark rearing in mice to investigate how light affects glucose and lipid homeostasis independent of disruptions in circadian rhythms. We show that 6-8 weeks of dark rearing from birth causes profound defects in glucose and lipid metabolism, including increased insulin secretion, depressed glucagon release, impaired hepatic gluconeogenesis and decreased lipolysis, all of which are consistent with attenuated sympathetic neuron activity. Indeed, sympathetic responses to cold exposure are attenuated in dark-reared animals. Metabolic defects in dark-reared animals are exacerbated with age, with older mice (>6 months) showing pronounced glucose intolerance, insulin resistance, increased body weight and fat mass, which were partially improved by exposure to 5 weeks of a regular light-dark cycle. Together, these results suggest that light exerts long-term effects on glucose and lipid metabolism via the sympathetic nervous system in a circadian-independent manner.

## Results

### Dark rearing results in enhanced glucose-stimulated insulin secretion in male mice

To investigate the chronic effects of light on glucose metabolism, we reared C57BL/6 mice in constant darkness (DD) from postnatal day 0 (P0) compared to control animals reared in a 12:12 light-dark (LD) cycle (**Fig. 1A**). We chose to raise the mice in constant darkness from birth, given previous studies that early life perturbations of environmental conditions result in pronounced effects on metabolism later on in life (*27, 28*). Mice were fed *ad libitum* with normal chow diet. We found that mice reared in DD maintained their endogenous circadian rhythms as assessed by locomotor activity patterns (**Fig. 1B**), although these mice had a slightly lengthened period. These results were similar to that observed in mice that were enucleated from birth (*29*). Neither the total daily locomotor activity nor total daily food intake were significantly different between animals raised in DD versus LD conditions (**Fig. 1, C and D**), although mice raised in DD tend to be more active during their subjective light phase compared to mice raised in the LD cycle (**fig. S1A**).

**Fig. 1.**
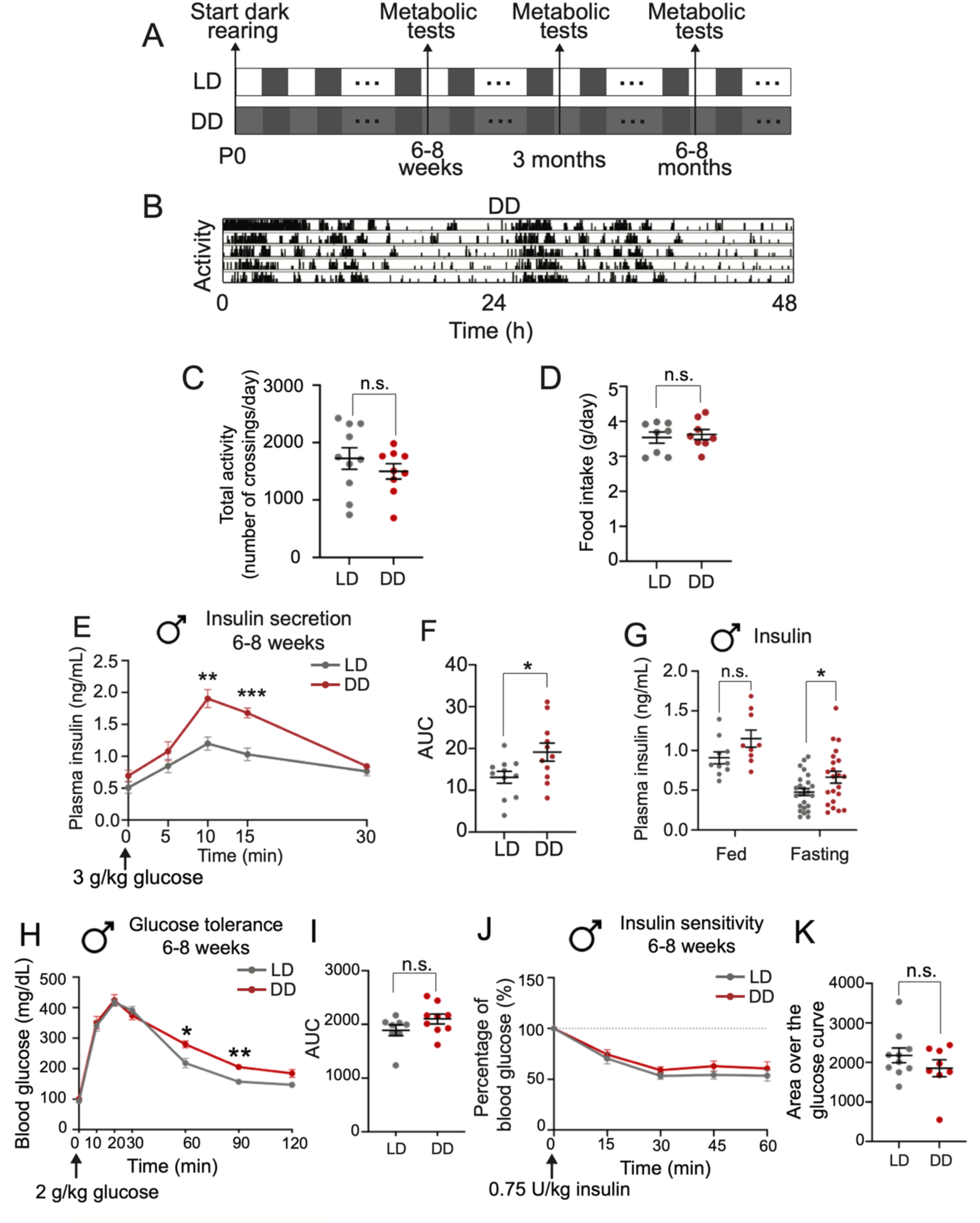
Dark rearing from birth results in increased insulin secretion in male mice at 6-8 weeks. **(A)** Schematic showing the experimental paradigm for animals reared in constant darkness (DD) from birth (Postnatal day 0, P0), compared to control animals reared in a 12:12 light-dark (LD) cycle. White, subjective day for control mice; dark grey, subjective night; light grey, subjective day for dark-reared mice. Metabolic tests were performed at 6-8 weeks, 3 months and 6-8 months after birth. **(B)** Representative actogram showing intact endogenous circadian rhythms in dark-reared mice, as assessed by locomotor activity. Activity is counted by the times that animal crosses the monitor in every 15 seconds. **(C)** Total locomotor activity is comparable between LD and DD mice. The total activity of an animal is measured by the total number of infrared monitor crossings per day. Results are means ± s.e.m with n=10 for LD and 9 for DD mice. n.s, not significant, unpaired t-test. **(D)** Total food intake (g/day) is similar between LD and DD mice. Results are means ± s.e.m with n=8 for each group. n.s, not significant, unpaired t-test. **(E, F)** Glucose-stimulated insulin secretion (GSIS) test. 6-8 weeks of dark-rearing increases plasma insulin levels in male mice. Area under the curve (AUC) of **(E)** is shown in **(F)**. Data are as means ± s.e.m with n=11 for each group. *p<0.05; **p<0.01; ***p<0.001, two-way ANOVA, Sidak’s multiple comparisons tests for **(E)** and unpaired t-test for **(F)**. **(G)** Plasma insulin levels. Fasting plasma insulin levels are elevated in male DD mice at 6-8 weeks. Data are means ± s.e.m for n=10 LD and 9 DD animals in fed condition, n=25 LD and 22 DD animals in fasting condition. *p<0.05, n.s, not significant, unpaired t-test. **(H, I)** Glucose tolerance tests. Glucose tolerance is mildly impaired in male DD mice after 6-8 weeks of dark rearing. AUC of **(H)** is shown in **(I)**. Results are means ± s.e.m for n=8 LD and 9 DD mice. * p<0.05; **p<0.01, two-way ANOVA, Sidak’s multiple comparisons tests for **(H)** and n.s, not significant, unpaired t-test for **(I)**. **(J, K)** Insulin sensitivity tests. Insulin sensitivity is unaffected by dark rearing for 6-8 weeks. Data are represented as percentages (%) of the blood glucose level at time “0”. Area over the glucose curve of **(J)** is shown in **(K).** The 100% blood glucose level at time 0 is set as the baseline. Results are means ± s.e.m with n=10 LD and 8 DD mice. Two-way ANOVA, Sidak’s multiple comparisons tests for **(J)** and n.s, not significant, unpaired t-test for **(K)**.

To assess glucose metabolism, we performed a battery of metabolic tests, including glucose-stimulated insulin secretion (GSIS), which measures plasma insulin levels in response to an oral glucose administration (*30, 31*); glucose tolerance tests (GTT), which measures the clearance of an orally administered glucose bolus from circulation (*32*); and insulin tolerance tests (ITT), which measures insulin responsiveness of peripheral tissues (*33*). Metabolic tests were performed in adult animals at various ages, i.e., 6-8 weeks, 3, and 6-8 months after birth. Given that there are daily rhythms in pancreatic hormone secretion, glucose tolerance, and glucose production (*34*), we performed metabolic tests at the same circadian time (CT), determined by locomotor activity patterns, for animals reared in DD or LD. We found that dark-reared mice had significantly higher plasma insulin levels in response to an oral glucose administration (3 g/kg body weight, gavage) compared to mice raised in LD, at 6-8 weeks of age (**Fig. 1, E and F**). Further, dark-reared mice also showed significantly higher plasma insulin levels under fasting condition (**Fig. 1G**). Notably, the enhanced insulin secretion and higher fasting insulin levels were specific to male mice; female dark-reared mice did not have any significant differences from those in the LD condition (**fig. S1, B to D**). Despite the increased plasma insulin levels *in vivo* in male dark-reared mice, isolated islets from these mice showed normal insulin secretory responses to glucose and KCl stimulation **(fig. S1E)**, indicating that insulin hyper-secretion *in vivo* did not stem from intrinsic defects in the islets themselves. Further, in assessing glucose tolerance and insulin sensitivity, we observed a modest impairment in glucose tolerance in dark-reared animals at 6-8 weeks compared to LD mice (**Fig. 1, H and I**), while insulin sensitivity was not affected (**Fig. 1, J and K**).

Together, these results indicate that 6-8 weeks of dark rearing primarily results in insulin hypersecretion, specifically in male mice, without significantly affecting glucose tolerance or insulin sensitivity at this stage.

### Dark-reared mice have impaired glucagon secretion, gluconeogenesis, and lipolysis at 6-8 weeks

Pancreatic islets play a critical role in blood glucose regulation by secreting the hormones, insulin and glucagon, in response to rising and falling glucose levels, respectively (*20*). Given the enhanced insulin secretion in dark-reared mice, we asked if glucagon levels might be affected. We measured plasma glucagon levels in response to insulin-induced hypoglycemia (*35*). In contrast to insulin, circulating glucagon levels, stimulated by hypoglycemia, were significantly attenuated in DD compared to LD animals at 6-8 weeks of age (**Fig. 2, A and B)**. Thus, pancreatic hormone secretion is impaired in dark-reared mice.

**Fig. 2.**
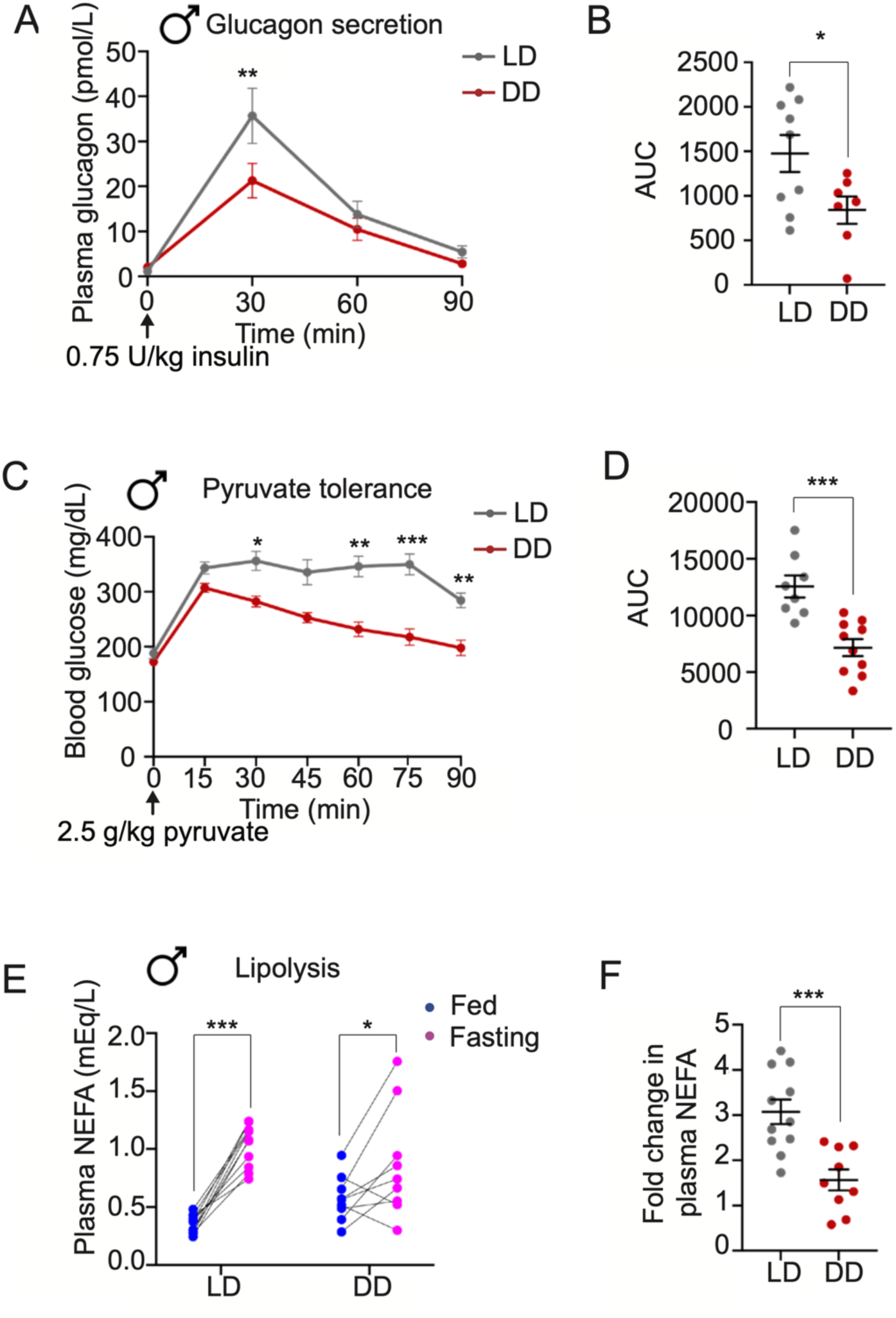
Dark reared mice show defects in glucagon secretion, gluconeogenesis, and plasma NEFA levels. **(A,B)** Glucagon secretion tests. Male DD mice show decreased glucagon secretion in response to insulin-induced hypoglycemia. Area under the curve (AUC) of **(A)** is shown in **(B)**. Results are means ± s.e.m with n=9 mice for LD and 7 for DD. *p<0.05, **p<0.01, two-way ANOVA, Sidak’s multiple comparisons tests for **(A)** and unpaired t-test for **(B)**. **(C, D)** Pyruvate tolerance tests show decreased gluconeogenesis in male DD mice after pyruvate injections. AUC of **(C)** is shown in **(D)**. Results are means ± s.e.m with n=8 mice for LD and 10 for DD. *p<0.05, **p<0.01, ***p<0.001, two-way ANOVA, Sidak’s multiple comparisons tests for **(C)** and unpaired t-test for **(D)**. **(E, F)** Lipolysis tests. Plasma non-esterified fatty acids (NEFA) levels measured by milliequivalents per liter (mEq/L) are elevated after fasting in both LD and DD animals **(E)**. However, the fold-increase in fasting plasma NEFA levels compared to the fed state is lower in DD animals **(F)**. NEFA fold change of each animal is calculated by dividing the fasting level by the fed level. Results are means ± s.e.m with n=11 mice for LD and 9 for DD. *p<0.05, ***p<0.001, two-way ANOVA, Sidak’s multiple comparisons tests for **(E)** and unpaired t-test for **(F)**.

The dynamic maintenance of blood glucose levels requires precise coordination of glucose uptake and endogenous glucose production. The liver plays a key role in regulating blood glucose homeostasis through the processes of *de novo* glucose production (gluconeogenesis) and glycogen breakdown (glycogenolysis). Since we observed insulin hypersecretion with dark-rearing, and insulin is known to suppress liver glucose production (*36*), we assessed gluconeogenesis using a pyruvate tolerance test, where pyruvate (2.5 g/kg pyruvate) delivered intra-peritoneally, is converted by the liver to glucose, which is then released into circulation (*37*). We found that circulating glucose levels in response to pyruvate administration were significantly decreased in DD compared to LD mice at 6-8 weeks of age (**Fig. 2, C and D**). These results suggest that hepatic gluconeogenesis is attenuated by dark rearing in mice.

As an additional measure of metabolic function, we assessed lipid metabolism by measuring plasma non-esterified fatty acids (NEFA), which are produced by lipolysis of triglycerides in white adipose tissue (WAT). Under fasting conditions, plasma NEFA levels rise rapidly, and this process is tightly regulated, with insulin and sympathetic nerve-derived Norepinephrine (NE) signaling being two major contributors (*38, 39*). We found that plasma NEFA increased both in mice reared in LD and DD after an overnight fast, when assessed at 6-8 weeks of age (**Fig. 2E**). However, the fold-increase in plasma NEFA triggered by fasting was significantly lower in mice reared in DD compared to LD conditions (**Fig. 2F**). These results suggest that 6-8 weeks of dark rearing results in attenuated WAT-mediated lipolysis in mice.

Similar to our results with insulin hyper-secretion, defective glucagon secretion, gluconeogenesis, and lipolysis at 6-8 weeks were specific to male mice; female dark-reared mice did not have any significant differences from those in the LD condition in all of these assessed metabolic functions (**fig. S2, A to F**).

### Sympathetic responses are reduced in dark-reared mice

The sympathetic nervous system is a key contributor to the control of glucose homeostasis (*20, 22*). Specifically, sympathetic nerve-derived norepinephrine (NE) signaling elevates circulating glucose levels by decreasing pancreatic insulin secretion and peripheral insulin sensitivity, increasing pancreatic glucagon secretion, promoting glycogen breakdown, and *de novo* glucose synthesis (gluconeogenesis) in the liver (*20–22*). Further, sympathetic activity also regulates energy balance, in part, by triggering lipolysis in white adipose tissue (*23*). Our findings that dark rearing results in increased insulin secretion, depressed glucagon release, reduced hepatic gluconeogenesis and WAT lipolysis at 6-8 weeks, together, suggested that sympathetic activity is suppressed in the absence of light input.

To directly address if sympathetic neuron activity is decreased in dark-reared mice, we performed immunostaining for c-Fos, an immediate early transcription factor, that serves as a robust reporter of neuronal activity (*40*). We chose to focus on the celiac-superior mesenteric ganglia complex (CG-SMG), since several metabolic organs, including the pancreas, liver, and gastrointestinal tract are predominantly innervated by post-ganglionic neurons that lie within this ganglion (*41–44*). At room temperature, we found that the number of c-Fos-positive sympathetic neurons in CG-SMG complex were not significantly different between LD and DD mice (**Fig. 3, A, B, and E**). However, activation of the sympathetic nervous system by a brief (1 hr) cold exposure at 4°C resulted in a robust increase in the number of c-Fos-positive neurons in ganglia in LD, but not in DD, animals (**Fig. 3, C to E**). These results suggest that sympathetic neuron activity, evoked in response to cold exposure, is attenuated in dark-reared animals. Of note, LD animals exposed to a 1 hr cold stimulus in the dark also showed a significant increase in c-Fos positive neurons (**fig. S3, A to C**), suggesting that the acute responses in sympathetic neuron activity is induced by cold exposure, and is not due to any effects elicited by performing experiments under light versus dark conditions.

**Fig. 3.**
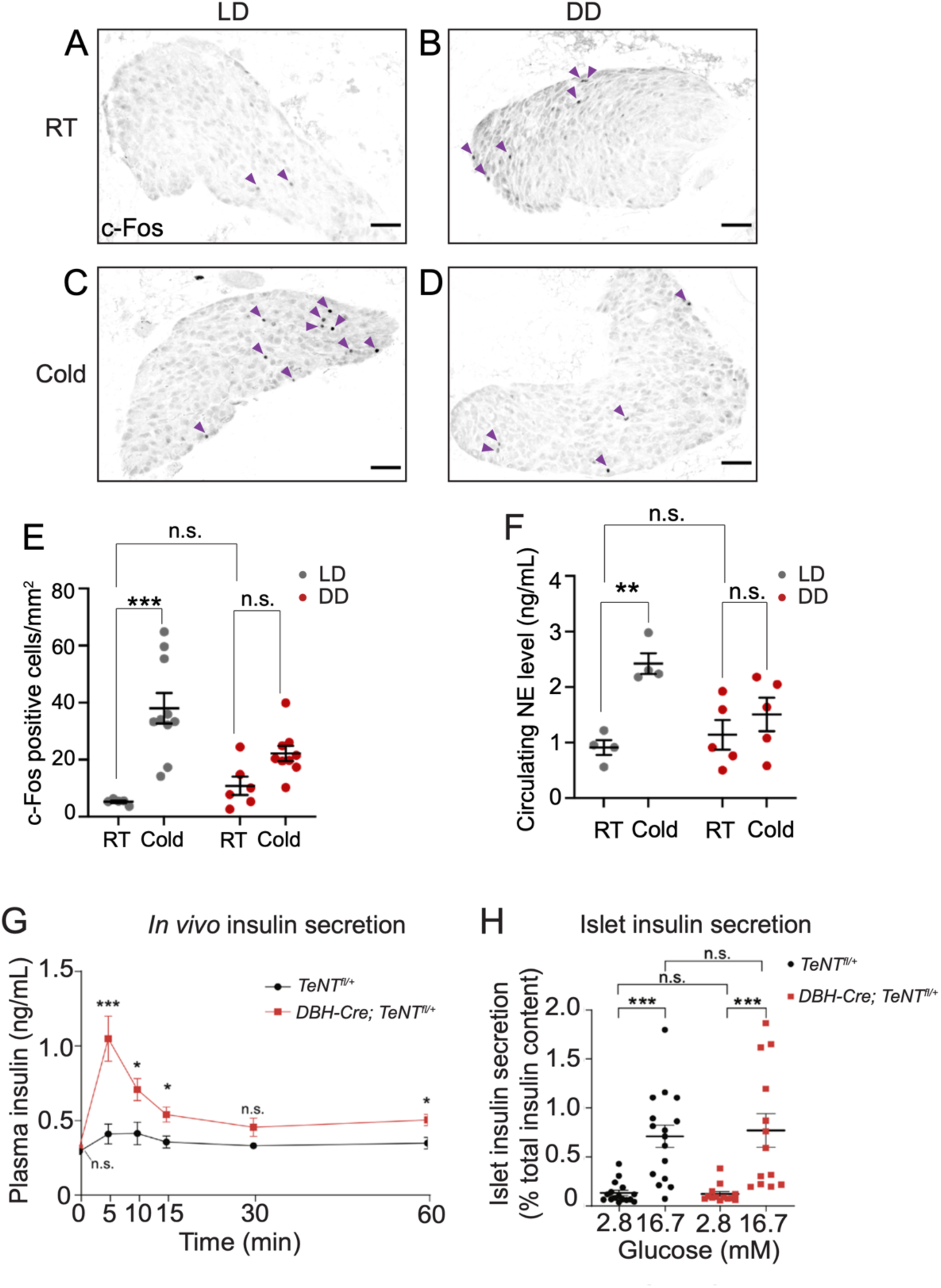
Sympathetic responses are attenuated in dark-reared mice. **(A-D)** c-Fos immunostaining in the celiac-superior mesenteric ganglion complex (CG-SMG) of LD and DD mice. RT, room temperature; Cold, 4°C for 1 hr. Scale bars; 100 μm. **(E)** Quantification of c-Fos-positive sympathetic neurons in CG-SMG from LD and DD mice at RT and in response to cold exposure (4°C, 1 hr). The numbers of c-Fos-positive neurons are similar between 6–8-week-old mice raised in LD and DD at RT. Cold exposure significantly increases number of c-Fos-positive neurons in mice reared in LD, but not, in DD. Data are presented as means ± s.e.m with n= 5 mice for LD and 6 for DD at RT; n=10 mice for LD and 9 for DD under cold exposure. ** p<0.01, n.s, not significant, two-way ANOVA, Sidak’s multiple comparisons tests. **(F)** Circulating norepinephrine (NE) levels of LD and DD mice. Cold exposure (4°C, 2 hr) significantly increases circulating NE in LD, but not DD, animals. Basal NE levels at room temperature are similar between LD and DD mice. Data are means ± s.e.m for n=4 mice for LD and 5 mice for DD. **p<0.01 n.s, not significant, two-way ANOVA, Sidak’s multiple comparisons tests. **(G)** Glucose-stimulated insulin secretion (GSIS) tests in DBH-Cre;TeNT^fl/+^ mice. Plasma insulin levels are elevated in *DBH-Cre;TeNT^fl/+^* mice compared to litter-mate controls (*TeNT^fl/+^*) at 6-8 weeks of age. Results are means ± s.e.m with n=6 control and 4 mutant mice. *p<0.05, ***p<0.001, n.s, not significant, unpaired t-test. **(H)** Basal and glucose-stimulated insulin secretion in isolated islets. Secreted insulin normalized to total insulin content is similar between islets isolated from *DBH-Cre;TeNT^fl/+^* mice and litter-mate controls (*TeNT^fl/+^*). Results are means ± s.e.m for islets isolated from n= 7 control and 6 mutant mice. ***p<0.001, n.s, not significant, unpaired t-test.

To further address the effects of dark rearing on sympathetic neurotransmission, we measured circulating levels of the neurotransmitter NE in LD and DD mice kept at room temperature or exposed to 4°C for 2 hr. Plasma NE levels are thought to be primarily derived from sympathetic nerves, although a smaller contribution comes from secretion from adrenal chromaffin cells (*45*). At room temperature, circulating NE levels were similar between LD and DD animals (**Fig. 3F**). In LD animals, cold exposure at 4°C significantly increased plasma NE compared to room temperature (**Fig. 3F**), consistent with elevated sympathetic activity. However, in DD animals, the brief cold exposure failed to enhance plasma NE (**Fig. 3F**), similar to the effects observed in c-Fos immunoreactivity in the ganglia. Together, these results suggest that dark rearing results in attenuated sympathetic responses in mice.

Given the blunted neuronal responses in DD animals, we asked if sympathetic axon innervation of target organs was altered by dark-rearing. Whole-organ immunostaining with the sympathetic neuron marker, Tyrosine Hydroxylase, in optically-cleared peripheral tissues, including the pancreas and kidney, followed by light sheet microscopy showed that axon innervation patterns were similar between LD and DD animals (**fig. S3, D and E**), suggesting that functional, rather than structural, defects underlie the decreased sympathetic responses elicited by dark rearing.

If reduced sympathetic activity contributes to defective glucose homeostasis in dark-reared mice, then silencing sympathetic activity should elicit similar metabolic phenotypes under a regular LD cycle. To test this prediction, we generated mice (*DBH-Cre;TeNT^fl/+^* mice), where sympathetic neurotransmission was blocked by expression of the tetanus toxin light chain subunit (*46*). We found that *DBH-Cre;TeNT^fl/+^*mice, that were 6-8 weeks old, raised in a regular LD cycle exhibit a striking increase in plasma insulin levels in response to an oral glucose administration (**Fig. 3G**), similar to our observations in dark-reared animals. Despite increased insulin secretion *in vivo*, isolated islets from *DBH-Cre;TeNT^fl/+^* mice showed normal insulin secretion to glucose stimulation, similar to that in dark-reared mice (**Fig. 3H**), indicating that insulin hyper-secretion *in vivo* does not stem from intrinsic defects in islets.

Together, these results suggest attenuated sympathetic neuron activity as a mechanism that contributes to metabolic defects in the absence of chronic light input.

### Metabolic defects in dark-reared mice are exacerbated with age

Most metabolic diseases are chronic and exacerbated with aging (*47*). Thus, we assessed metabolic function in dark-reared animals at older ages, specifically, at 3 months and 6-8 months. We found that after 3 months of dark rearing, mice developed insulin resistance, assessed by an insulin tolerance test (ITT) (**Fig. 4, A and B**), which was maintained in dark-reared animals at 6 months of age (**fig. S4, A and B**). In assessing glucose tolerance, dark-reared mice developed prominent glucose intolerance by 6 months of age (**Fig. 4, C and D**), although this parameter was only modestly impaired in DD animals at 6-8 weeks (see **Fig. 1, H and I**) and at 3 months (**fig. S4, C and D**).

**Fig. 4.**
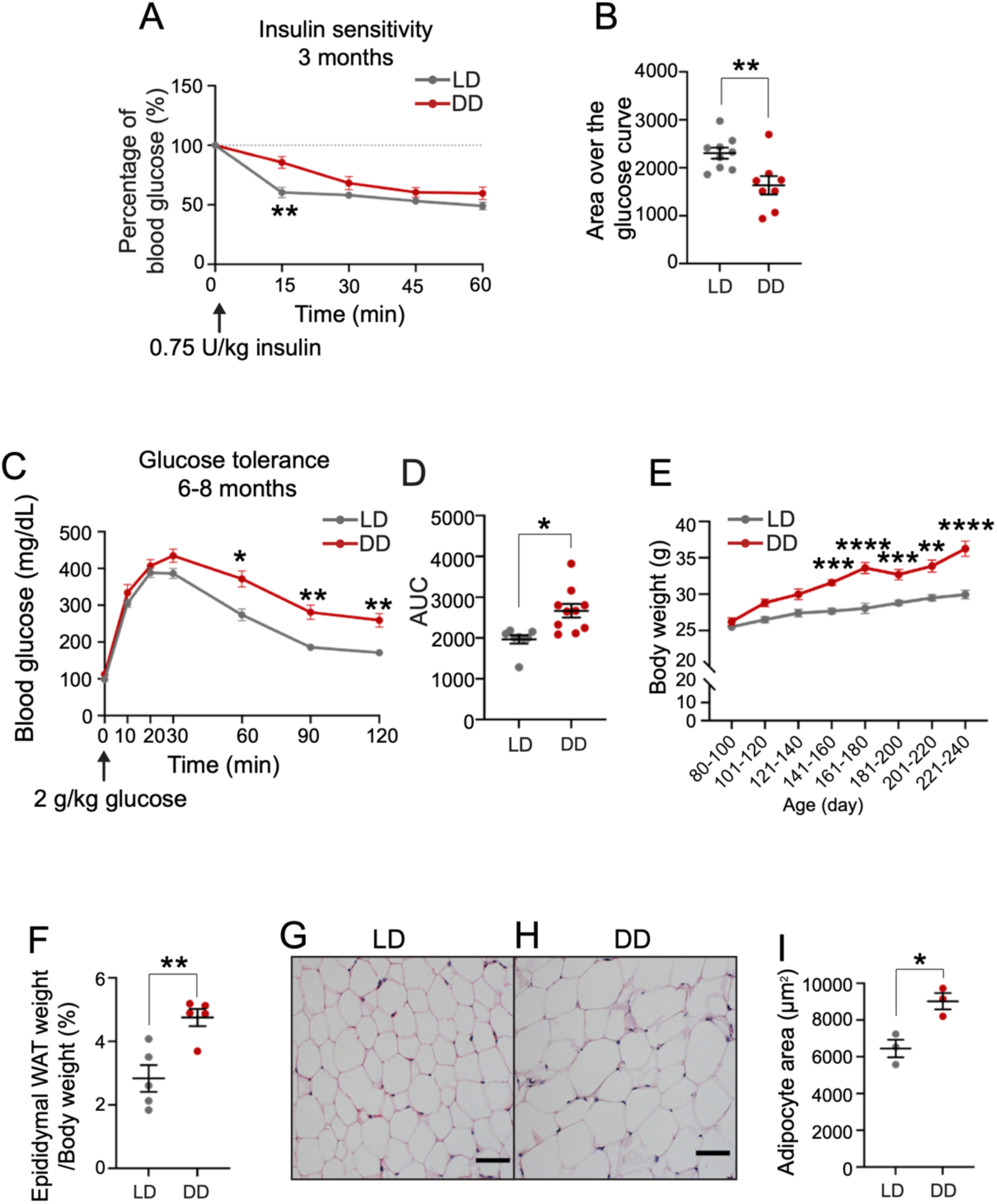
Dark-reared animals develop insulin resistance, glucose intolerance, and gain in body weight with age. **(A,B)** Insulin sensitivity tests. Dark-reared male mice become insulin resistant by 3 months. Data are represented as percentages (%) of the blood glucose level at time “0”. Area over the glucose curve of **(A)** is shown in **(B).** The 100% blood glucose level at time 0 is set as the baseline. Results are means ± s.e.m for n=9 mice for LD and 8 for DD. **p<0.01, two-way ANOVA, Sidak’s multiple comparisons tests for **(A)** and unpaired t-test for **(B)**. **(C, D)** Glucose tolerance tests. Dark-reared male mice develop glucose intolerance at 6-8 months. Area under the curve (AUC) of **(C)** is shown in **(D)**. Results are means ± s.e.m with n=8 mice for LD and 10 for DD. *p<0.05; **p<0.01, two-way ANOVA, Sidak’s multiple comparisons tests for **(C)** and unpaired t-test for **(D)**. **(E)** Body weight of LD and DD mice. Dark rearing causes increased body weight with age in male mice. Results are means ± s.e.m with n=6-32 mice for LD and 8-20 mice for DD. ** p<0.01; *** p<0.001; **** p<0.0001, two-way ANOVA, Sidak’s multiple comparisons tests. **(F)** Weight of epididymal white adipose tissue (WAT) of LD and DD mice. Dark-reared male mice accumulate WAT compared to mice raised in LD cycle at 6-8 months measured by percentage of the epididymal WAT weight normalized by the body weight. Results are means ± s.e.m for n=5 each group. ** p<0.01, unpaired t-test. **(G, H)** H&E staining shows that adipocytes are larger in DD animals **(H)** compared to LD **(G)** at 6-8 months. Scale bar: 100 μm. **(I)** Quantification of average adipocyte size (μm^2^) in LD versus DD animals. Results are means ± s.e.m for n=3 mice for each group. * p<0.05, unpaired t-test.

Notably, dark-reared animals gained in body weight with age (**Fig. 4E**). These animals showed significantly increased body weight starting at 5 months of age compared to animals raised in LD (**Fig. 4E**), and without differences in total locomotor activity or food intake between the two groups (see **Fig. 1, C and D**). Consistent with the increased body weight, we found that dark-reared animals gained in fat mass, as assessed by an increase in epididymal white adipose tissue (**Fig. 4F**). We found that epididymal WAT contributed to about 4.8% of body weight in mice reared in DD, compared to only 2.8% of body weight in animals reared in LD (**Fig. 4F**). Further, Hematoxylin and Eosin staining showed significant hypertrophy of adipocytes in dark-reared animals (**Fig. 4, G to I**).

Thus, dark-reared animals show an exacerbation of defects in glucose metabolism with age, with 6-8 month-old mice showing pronounced glucose intolerance, insulin resistance, and increased body weight and fat mass.

### Exposure to a regular LD cycle partially alleviates metabolic defects in dark-reared mice

We next asked if metabolic abnormalities caused by long-term dark rearing can be reversed by light exposure. Mice that were dark-reared for 10 months were placed under a regular 12:12-hour LD cycle for 5 weeks. Metabolic functions were tested before and after exposure to LD cycle (**Fig. 5A**). We found that body weights of these mice were not significantly different after exposure to a regular LD cycle (**Fig. 5B**). However, we observed a pronounced drop in fasting glucose levels in animals upon exposure to the LD cycle (**Fig. 5C**). Further, 5 weeks in the regular LD cycle improved glucose tolerance and insulin sensitivity in animals that had been dark-reared for 10 months (**Fig. 5, D to G**).

**Fig. 5.**
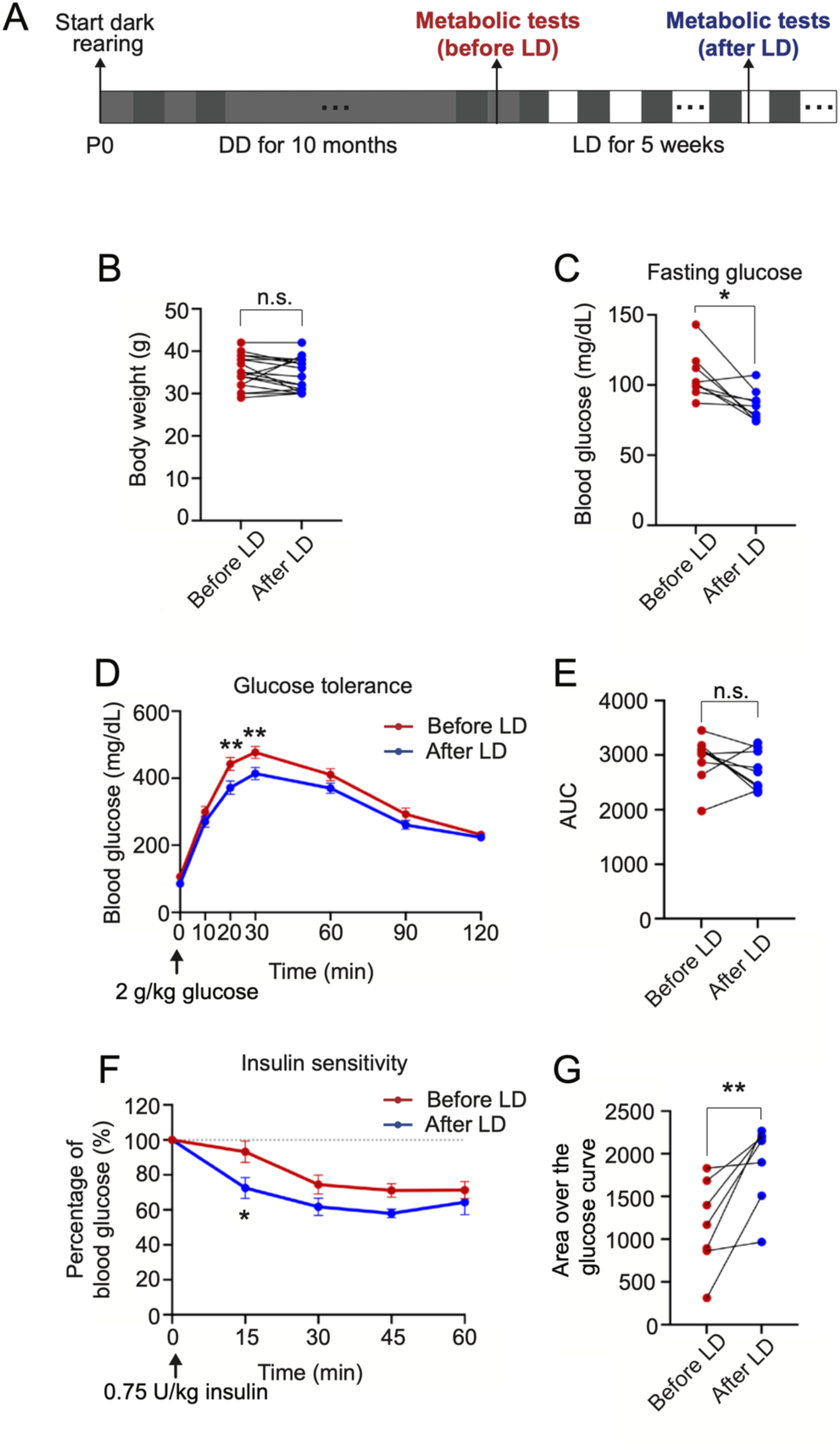
Exposing dark-reared animals to a regular light/dark cycle partially alleviates metabolic defects. **(A)** Schematic diagram showing that mice were reared in DD from P0 for 10 months and then moved to a regular 12:12 LD light condition for another 5 weeks. Metabolic tests were performed immediately before, and after, the 5 weeks of LD exposure. Light grey, subjective day for DD phase; dark grey, subjective night; white, subjective day for LD phase. **(B)** Body weights were unaffected by 5 weeks of LD exposure in animals that were reared in DD for 10 months. Results are means ± s.e.m for n=17 mice. n.s, not significant, paired t-test. **(C)** Fasting glucose levels were reduced after 5 weeks of LD exposure in mice dark-reared for 10 months. Results are means ± s.e.m for n=9 mice. *p<0.05, paired t-test. **(D, E)** Glucose tolerance was improved after 5 weeks of exposure to a regular LD cycle. Area under the curve (AUC) of **(D)** is shown in **(E)**. Results are means ± s.e.m for n=9 mice. **p<0.01; two-way ANOVA, Sidak’s multiple comparisons tests for **(D)** and paired t-test for **(E)**. **(F, G)** Insulin sensitivity was improved by 5 weeks of LD exposure. Data are represented as percentages (%) of the blood glucose level at time “0”. Area over the glucose curve of **(F)** is shown in **(G).** The 100% blood glucose level at time 0 is set as the baseline. Results are means ± s.e.m for n=7 mice. * p<0.05; **p<0.01, two-way ANOVA, Sidak’s multiple comparisons tests for **(F)** and unpaired t-test for **(G)**.

Together, these results indicate that 5 weeks of exposure to a regular 12:12 LD cycle is capable of partially alleviating metabolic defects caused by long-term light deprivation, and reveal the plasticity of light-mediated effects on glucose metabolism.

## Discussion

Classically, circadian disruption caused by irregular light conditions has been thought to be the primary reason for perturbations in glucose metabolism and energy balance in animal models and humans (*3, 48*). Using dark reared mice that maintain intact endogenous circadian rhythms, our studies reveal a non-canonical pathway by which light/dark conditions affect metabolism. In this study, we show that long-term light deprivation from birth, in the context of undisturbed circadian rhythms, results in attenuated sympathetic activity and metabolic defects in mice, especially in males. In dark-reared mice, tissue-specific metabolic functions regulated by the sympathetic nervous system, specifically, pancreatic hormone secretion, hepatic gluconeogenesis, and white adipose tissue lipolysis, were impaired by 6-8 weeks, whereas phenotypes related to overall whole-body metabolism, including insulin resistance, glucose tolerance, and increased body weight, emerged later with age. Strikingly, the metabolic defects were partially mitigated by exposing animals to a regular 12:12 hour light-dark cycle for 5 weeks. Together, our findings indicate that long-term light-mediated modulation of the sympathetic nervous system is essential for metabolic function in mice, and that regular light exposure can partially reverse deficits elicited by chronic light deprivation, revealing the plasticity of light-mediated effects in adult animals.

To date, the handful of studies that have shown a direct circadian-independent effect of light on peripheral tissue physiology have relied exclusively on acute light stimulation (∼ 2 hr) (*10, 11*). Our results of metabolic dysfunction in animals raised in the dark from birth provide insight into chronic effects of light on metabolic physiology that occur in a circadian-independent manner. We found that dark rearing of mice from birth resulted in metabolic defects that manifested at different stages across a long duration in the life of animals, and were exacerbated with age. Specifically, we observed impaired pancreatic hormone secretion, hepatic gluconeogenesis, and white adipose tissue lipolysis in young adult mice after 6-8 weeks of dark rearing, while older animals developed insulin resistance by 3 months, followed by severe glucose intolerance and increased body weight at 6-8 months. It appears paradoxical, at first glance, that dark-reared animals first showed insulin hyper-secretion in early life before progressing to insulin resistance and glucose intolerance at older ages. However, our findings align with recent unexpected evidence in humans and in animal models that insulin hyper-secretion is a triggering event in the development of type 2 diabetes, likely through promoting insulin resistance (*49–52*). While it is well-known that visual impairment is a long-term consequence of diabetes (*53*), whether humans with visual impairment are at increased risk for developing type 2 diabetes remains unclear. Nevertheless, our findings underscore the importance of exposure to a regular light/dark cycle on metabolic health. Of note, we specifically observed metabolic defects in male mice with dark rearing. In general, multiple studies have noted that in both humans and mice, females are resistant to metabolic challenges (*54, 55*), likely due to the protective effects of estrogens (*56*). Further, we observed metabolic defects in dark-reared animals that were maintained on a standard chow diet. It will be of interest in future studies to assess if the metabolic phenotypes observed with dark rearing are exacerbated with raising animals on high-fat or high-sugar diets.

Our results suggest that light signals impinging on the retina are relayed to peripheral metabolic organs via the sympathetic nervous system. Since their discovery in 2002 (*6, 57*), ipRGCs are now known to be critical for relaying photic information to control several non-image functions, including circadian photoentrainment, sleep, mood, and cognitive functions (*1, 7, 8*). In addition to these well-studied functions, limited studies suggest that ipRGCs can also influence the functions of peripheral tissues, including thermogenesis in brown adipose tissue and the proliferation of hair follicle stem cells in the skin, by acting via the sympathetic nervous system (*10–12*). However, the neuroanatomical connections between ipRGCs and sympathetic ganglia remain undefined. In particular, the neural connections that allow light information to be relayed to peripheral glucose-regulatory organs via the sympathetic nervous system remain to be identified. ipRGCs directly innervate several hypothalamic brain regions including the SCN, the supraoptic nucleus (SON), and lateral hypothalamic area (LHA), as well as form secondary connections downstream of the SON with the paraventricular nucleus (PVN), that are known to be involved in blood glucose regulation, pancreatic hormone secretion, and lipid metabolism (*58*). The PVN is a region of particular interest; although the PVN does not receive direct ipRGC input, it has reciprocal innervation with SCN (*59, 60*), which is directly innervated by ipRGCs (*58, 60*). The PVN also receives innervation from the SON, which receives direct ipRGC input (*10, 60*). The PVN is well-positioned to relay CNS outflow to the sympathetic nervous system, since it contains pre-autonomic neuronal populations that either directly or indirectly connect to pre-ganglionic sympathetic neurons in the spinal cord (*10, 61–63*). Future tracing analyses to identify the neuroanatomical pathways connecting ipRGCs to peripheral sympathetic neurons innervating metabolic tissues will be critical to advance the overall understanding of how light influences glucose metabolism.

Remarkably, we found that five weeks of exposure to a regular light/dark cycle partially improved glucose intolerance and insulin resistance in animals that were dark-reared for 10 months, suggesting that the detrimental effects of chronic light deprivation on glucose metabolism are reversible, even in older animals. In dark-reared animals, we found that metabolic defects manifested without any alterations in neuronal innervation to peripheral organs, suggesting that functional, rather than structural, defects in neurons underlie the metabolic defects in the absence of light input. Further, isolated islets from dark-reared mice were capable of responding normally to elevated glucose stimulus *in vitro*, suggesting light deprivation did not elicit intrinsic defects in glucose-regulatory peripheral tissues and/or organs themselves. These factors may explain why re-exposing the animals to a regular light-dark cycle results in a relatively rapid reversal of metabolic dysfunction.

Light therapy has been widely used to treat mood disorders, including seasonal affective disorder, depression, and bipolar disorder (*5, 64*). In addition to improving psychiatric status, epidemiological studies have shown that light stimulation also exerts beneficial effects on glucose control and whole-body physiology in individuals (*5, 65, 66*), including in individuals with type 2 diabetes (*67, 68*). It has been assumed that the beneficial effects of light therapy on metabolic functions are primarily mediated by re-adjusting a misaligned SCN and peripheral clocks (*5*). Our findings provide insight into circadian-independent mechanisms by which light modulation influences whole-body physiology. A complete understanding of all the neural mechanisms by which light influences metabolic physiology is critical to inform new strategies for treating disorders such as type 2 diabetes and obesity that are linked to aberrant light/dark conditions.

## Materials and Methods

### Animals

All animal care and experimental procedures were conducted in accordance with the Johns Hopkins University Animal Care and Use Committee (ACUC) and NIH guidelines. All efforts were made to minimize the pain and number of animals used. Control animals were group housed in a standard 12:12 light-dark (LD) cycle, while light deprivation was done by rearing animals in a constant dark room from birth, with water and food provided *ad libitum*. The circadian time (CT) was used to determine the time for all experimental analyses with the start of active phase defined as CT12. LD or DD light conditions were maintained during all the experiments, except for experiments in Figure 5 where mice raised in constant darkness for 10 months were exposed to a regular 12:12 LD cycle for 5 weeks. Ages of mice are indicated in the figure legends and/or methods. The following mouse lines were used in this study: C57/B6J (JAX# 000664) mice were obtained from The Jackson Laboratory. *DBH-Cre;TeNT^fl/+^*mice were generated by crossing *DBH-Cre* mice (*69*) (a gift from Dr. Warren G. Tourtellotte Northwestern University) with *TeNT^fl/fl^* mice (*46*) (from Dr. Martyn Goulding, Salk Institute).

### Locomotor activity monitoring

Mouse locomotor activity was monitored by an infrared motion detector (STAR Life Sciences Corp) built into the mouse cages. VitalView® 6 software (STAR Life Sciences Corp) was used to measure activity levels, and a customized Python code was used to generate actograms to determine the start of the active phase, which was assigned as CT12 for DD animals. The activity level was calculated as the total number of crossings detected in every 15 seconds.

### Food intake measurement

Mice (6-10 months) were weighed and then individually housed for a total of 7 days. 100gm of food was given to each mouse on the first day. The weight of uneaten food was measured each day, and total food intake per day was calculated by averaging the intake food each day throughout the 7 days.

### Glucose-induced insulin secretion (GSIS) test

Mice at indicated ages were individually housed and fasted for 16 hr from CT9 to CT1 the next day, after which mice were given 3 g/kg glucose (Sigma-Aldrich) by oral gavage. Tail blood was collected at the indicated times in DTA-coated tubes, spun down at 3500 rpm for 15 min at 4°C, and plasma insulin levels were measured with an Ultrasensitive Insulin ELISA kit (Crystal Chem). Fed and fasting insulin levels were measured from blood samples before and after fasting.

### Glucose tolerance test

Mice at indicated ages were individually housed and fasted for 14-16 hr from CT9-11 to CT1 the next day. After fasting, mice were given 2 g/kg glucose (Sigma-Aldrich) by oral gavage, and tail blood (∼5μl) glucose levels were measured using OneTouch Ultra glucometer at the indicated times.

### Insulin sensitivity test

Mice at indicated ages were individually housed and fasted for 2 hr from CT2 to CT4 before being injected intraperitoneally (i.p.) with 0.75 U/kg of insulin (Novolin-R). Tail blood glucose levels (∼5 μl) were measured using a OneTouch Ultra glucometer at indicated times as previously described (*70*).

### Glucagon secretion

Mice (6-8 weeks) were individually housed and fasted 2 hr from CT 2 to CT 4. insulin (Novolin-R) was i.p. injected at 0.75 U/kg. Tail blood (∼20 μl) was collected at 0, 30, 60 and 90 min after insulin injection in DTA-coated tubes, spun down at 3.5 rcf for 15 min, and plasma was collected to PCR tubes containing aprotinin (Sigma-Aldrich) and stored in −80 °C. Plasma glucagon levels were measured with a commercial mouse glucagon ELISA kit (Mercodia) according to manufacturer’s protocol.

### Pyruvate tolerance test

Mice (6-8 weeks) were individually housed and fasted 6 hr from CT 2 to CT8. After fasting, mice were given 2.5 g/kg pyruvate (Sigma-Aldrich) by i.p. injection, and tail blood (∼5 μl) glucose levels were measured using OneTouch Ultra glucometer at indicated time points.

### Lipolysis test (plasma NEFA levels)

Mice (6-8 weeks) were individually housed and fasted 16 hr from CT9 to CT1 the next day. Tailed blood (∼ 40ul) was collected before and after fasting in DTA-coated tubes, spun down at 3.5 rcf for 15 min at 4°C and stored at −80 °C, and plasma NEFA levels were measured with FUJIFILM Medical Systems USA Hr Series Nefa-Hr (2) kit (Fisher Scientific).

### Mouse islet isolations, *in vitro* insulin secretion and insulin content measurement

Pancreatic islets were isolated from mice at 6-8 weeks of age as previously described (*71*). Briefly, animals were killed by cervical dislocation and collagenase solution (Collagenase P [Roche], 0.375 mg/mL in HBSS) was injected through the bile duct. Pancreata were dissected and digested with 3 ml collagenase solution at 37°C for 10 min. Digested pancreata were washed with HBSS containing 0.1% BSA and subjected to HBSS/ histopaque (6:5 Histopaque 1119:1077; Sigma) discontinuous density gradient centrifugation. The islet layer found at the interface was collected, and islets were handpicked under an inverted microscope. For insulin secretion in isolated islets, harvested islets were allowed to recover overnight in RPMI-1640 media containing 10% fetal bovine serum, and 5 U/L penicillin/streptomycin (Invitrogen). Islets were washed with Krebs-Ringer HEPES buffer (KRBH) containing 2.8 mM glucose and pre-incubated in the same fresh buffer for 1 hr. After pre-incubation, groups of similarly sized 8-10 islets were handpicked into a 24-well plate and allowed to incubate for 30 min in KRBH containing 2.8 mM glucose. Islets were transferred to KRBH containing 16.7 mM glucose and incubated for 30 min, and then transferred to 30 mM potassium chloride and incubated for 30min. After each round of incubation, the supernatant fractions were collected. After the last incubation, islets were lysed with cold acid ethanol. Insulin content in both supernatant and islet fractions was determined by insulin ELISA kits (Crystal Chem). The secretion was calculated by the percentage of insulin in the fraction versus the total insulin levels in cell lysates plus supernatants.

### White adipose tissue mass measurement and Hematoxylin and Eosin (H&E) staining

Mice (8-10 months) were weighted and euthanized with CO_2_ inhalation. Epidydimal white adipose tissue (WAT) was dissected and weighted. For H&E staining, epidydimal WAT was fixed in Bouin’s solution for 3 hr at room temperature (RT), washed with 70% ethanol and kept overnight in 70% ethanol at RT. Tissues were then dehydrated with sequential treatment of 95% ethanol, 100% ethanol, and xylene. Tissues were embedded in paraffin and sectioned at 6 μm using a microtome. Tissue sections were then dried at 60°C for 1.5 hr. Tissue sections were deparaffinized using xylene, and rehydrated using 100%, 95%, 80%, 70% and deionized water. The tissues were then incubated in Hematoxylin Solution, Harris Modified (Sigma-Aldrich) and rinsed in water, followed by incubation in 1% lithium carbonate, deionized water, and 80% ethanol. Tissues were incubated in eosin Y solution (Sigma-Aldrich) and rinsed in 80%, 95%, and 100% ethanol and xylene. Tissues were air dried and mounted with Permount mounting media (Thermo Fisher Scientific). Images were taken under bright field using Zeiss upright microscope. Adipocyte areas were quantified using Image J (Fiji) software.

### c-Fos immunohistochemistry

Mice (8-10 weeks) were euthanized with CO_2_ inhalation. The CG-SMG tissue was dissected from mice that were kept at room temperature or exposed to cold (4°C, 1 hr) and fixed with 4% paraformaldehyde (PFA) overnight at 4 °C. Tissues were cryoprotected in 30% sucrose/PBS for 2 hr at 4 °C and embedded in OCT and stored at −80°C. 20 μm thick sections were cut using a cryostat. Sections were collected and washed 4 times in 0.3% Triton-X 100 in PBS (0.3% PBST) for 5 min each, incubated in 0.1 M glycine for 5 min, followed by 9 times in 0.3% PBST for 5 min each. Samples were incubated in blocking solution (5% goat serum in 0.3% PBST) for 1.5 hr at RT, followed by incubation with a polyclonal (Abcam) or monoclonal (Abcam) rabbit anti-c-Fos (1:1000) antibody diluted in blocking solution overnight at 4 °C. After washes, sections were incubated in Alexa-546 conjugated goat anti-rabbit secondary antibody (Thermo Fisher Scientific) (1:400) overnight at 4 °C. Samples were washed and mounted using Fluoromount (Sigma-Aldrich) and imaged using LSM 780 confocal scanning microscope. Number of c-Fos-positive cells were quantified by manual counting and the area of the ganglion section was calculated using Fiji software.

### Plasma norepinephrine (NE) measurement

Blood samples (300 μl) were drawn retro-orbitally from anesthetized mice that were either housed at room temperature or exposed to cold (4°C, 2 hr). Plasma was prepared by centrifuging blood at 3.5 rcf for 15 min at 4°C and stored at −80°C. 100 μl plasma was used for NE measurements using ELISA (NE High Sensitive ELISA Kit, Rocky Mountain Diagnostics) according to manufacturer’s protocol.

### FLASH wholemount immunostaining and tissue clearing

FLASH-based whole mount immunostaining and tissue clearing of pancreas and kidneys from 6–8-week-old mice was performed as previously described (*72*) with modification. Briefly, mice were transcardially perfused with 4%PFA. Pancreas and kidneys were dissected and post-fixed in 4%PFA overnight at 4°C, then washed with PBS and incubated in a borate-based antigen retrieval solution overnight at 54°C. Samples were then permeabilized in 0.2% Triton X-100 in PSB (0.2% PBST) for 3 hr and blocked by blocking solution (0.2% PBST/10%DMSO/6% Goat Serum) for 3 hr at RT and then incubated with rabbit-anti-TH (1:500) and mouse-anti-insulin (1:2000) in blocking solution for 5 days at RT. After washing 3 times with 0.2% PBST for 1 hr each and overnight, samples were incubated with goat-anti-rabbit/mouse secondary antibody (1:400) for 5 days at RT. Samples were then washed 3 times with 0.2% PBST for 1 hr each and overnight and dehydrated, in sequential, in 20%, 40%, 60%, 80% and 100% methanol, followed by 25%, 50%, 75%, 100% Benzyl Alcohol/Benzyl Benzoate (BABB) in methanol. Samples were stored in 100% BABB until imaging. Tissues were imaged using LSM 780 confocal microscope.

### Quantification and statistical analyses

For practical reasons, all metabolic analyses, immunostaining, and adipocyte analysis were performed in semi-blinded manners. For LD and DD mice, the experimenter only kept track of LD versus DD conditions during the sample collecting part and did the measurements blindly. For *DBH-Cre;TeNT^fl/+^* and littermate *TeNT^fl/+^* control mice, experiments were done blindly by only knowing the animal’s ear-tag number. All graphs and statistical analyses were performed using GraphPad Prism 9. Statistical significance was determined using unpaired two-tailed student t-tests for unpaired samples, paired two-tailed student t-tests for paired samples, and two-way ANOVA tests with Sidak’s multiple comparisons tests for more than one variable. All error bars are represented as the standard error of the mean (s.e.m).

## Acknowledgements

We thank the JHU Integrated Imaging Center for assistance with microscopy. This study was supported by NIH R01 awards, NS114478 and NS107342, to R.K, DC016065 and EY027202 to H.Z, and NIMH intramural research funds (ZIAMH002964) to S.H.

## Author Contributions

X.C., R.K., H.Z. and S.H. contributed to study design, and writing and editing the manuscript. X.C. conducted the majority of experiments and analyzed data. E.L. performed the GSIS assays in *DBH-Cre; TeNT^fl/+^* mice. M.H. assisted with the metabolic tests, and L.W.S contributed to analyses of sympathetic innervation.

## Declaration of Interests

The authors declare no competing interests.

## Materials and data availability

All transgenic mice generated in this study are available upon request. All data needed to evaluate the conclusions in the paper are present in the paper and/or the Supplementary Materials. Any additional information required to re-analyze the data reported in this paper is available from the lead contact upon request. This paper does not report original code.

---

**Fig. S1.**
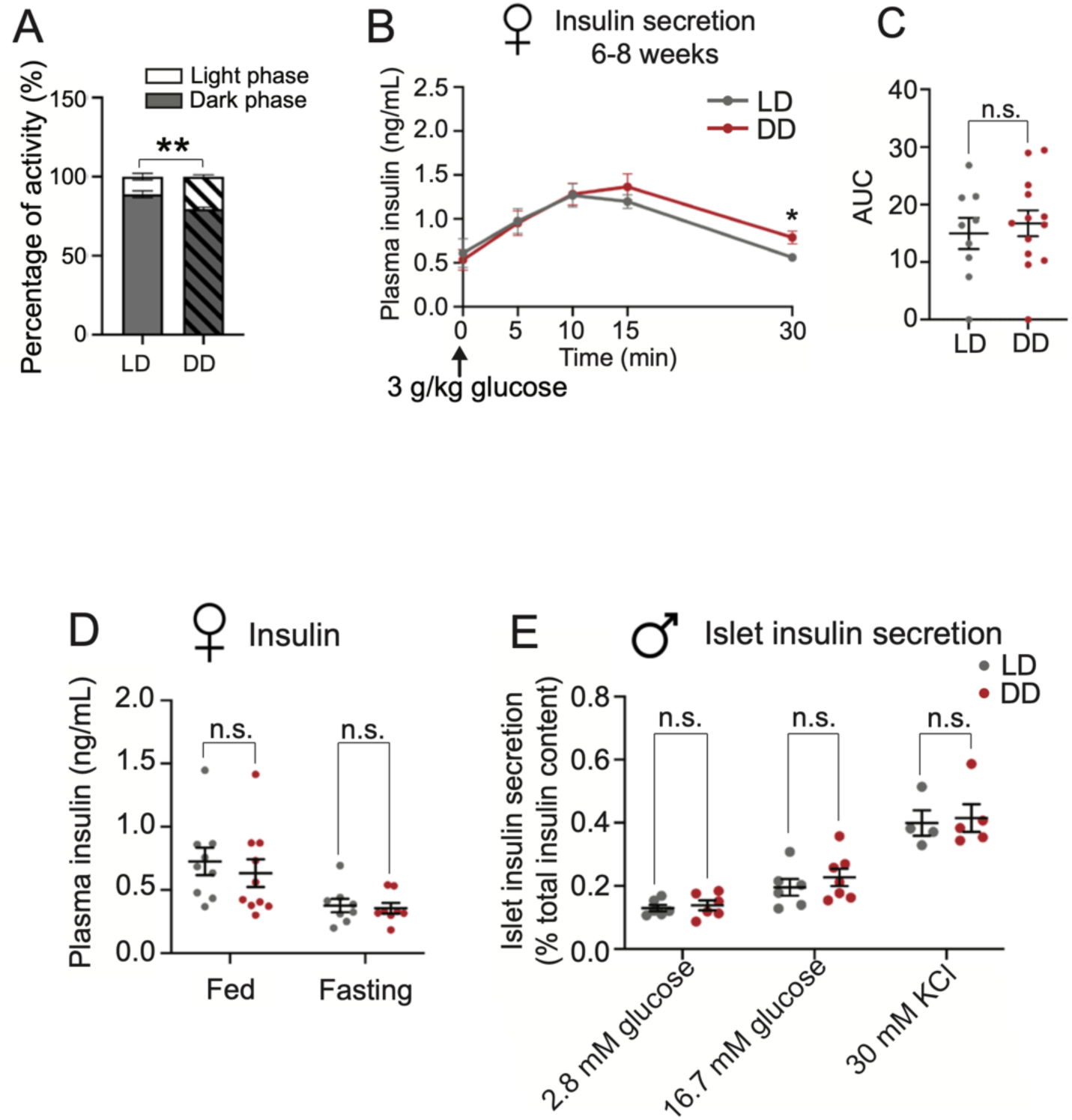
Dark rearing does not affect insulin secretion in female mice. **(A)** The amount of locomotor activity is differentially distributed between LD and DD mice through the day. Dark-reared mice have a higher portion of the activity during the subjective day compared to mice in LD. Results are means ± s.e.m with n=10 mice for LD and 9 for DD. **p<0.01, two-way ANOVA, Sidak’s multiple comparisons tests. **(B, C)** Glucose-stimulated insulin secretion (GSIS) tests. Plasma insulin levels are unaffected in female mice raised in DD. Data are means ± s.e.m with n=9 for LD and 13 for DD. *p<0.05, two-way ANOVA, Sidak’s multiple comparisons tests for **(B)** and n.s, not significant, unpaired t-test for **(C)**. **(D)** Plasma insulin levels. Female mice raised in DD and LD show similar fed and fasting insulin levels. Data are as means ± s.e.m with n=9 mice for LD and 10 for DD for fed insulin levels; n=8 mice per group for fasting insulin; n.s, not significant, unpaired t-test. **(E)** Isolated islets from male mice raised in DD have similar basal insulin secretion and secretory responses to high glucose (16.7 mM) or KCl (30 mM) as those from male mice raised in LD. Data are means ± s.e.m with isolated islets from n=4-7 mice for each group. n.s, not significant, two-way ANOVA, Sidak’s multiple comparisons tests.

**Fig. S2.**
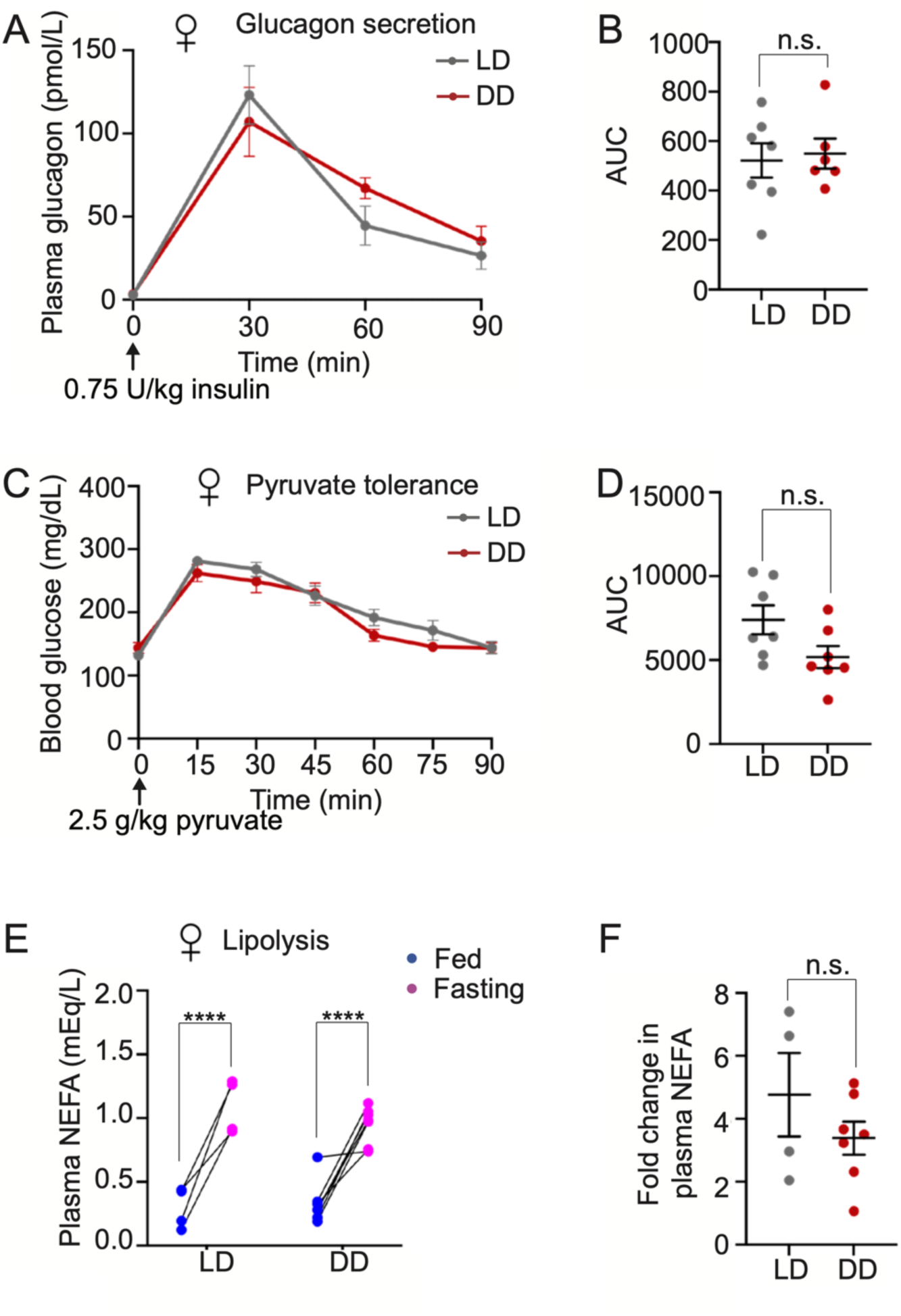
Dark rearing does not affect glucagon secretion, gluconeogenesis, or lipolysis in female mice. **(A,B)** Glucagon secretion tests. Glucagon secretion stimulated by insulin injection is similar between female mice reared in LD and DD at 6-8 weeks. Area under the curve (AUC) of **(A)** is shown in **(B)**. Results are means ± s.e.m with n=7 mice for LD and 6 for DD. Two-way ANOVA, Sidak’s multiple comparisons tests for **(A)** and n.s, not significant, unpaired t-test for **(B)**. **(C, D)** Pyruvate tolerance tests show that gluconeogenesis is unaffected in female mice reared in DD, at 6-8 weeks. AUC of **(C)** is shown in **(D)**. Results are means ± s.e.m with n=7 mice for each group. Two-way ANOVA, Sidak’s multiple comparisons tests for **(C)** and n.s, not significant, unpaired t-test for **(D)**. **(E, F)** Lipolysis tests. Plasma non-esterified fatty acids (NEFA) levels measured by milliequivalents per liter (mEq/L) are elevated after fasting in both LD and DD animals **(E)**. The fold-increase in plasma NEFA levels after fasting is similar in LD and DD animals **(F)**. NEFA fold change of each animal is calculated by dividing the fasting level by the fed level. Results are means ± s.e.m with n=4 mice for LD and 7 for DD. **** p<0.0001, two-way ANOVA, Sidak’s multiple comparisons tests for **(E)** and n.s, not significant, unpaired t-test for **(F)**.

**Fig. S3.**
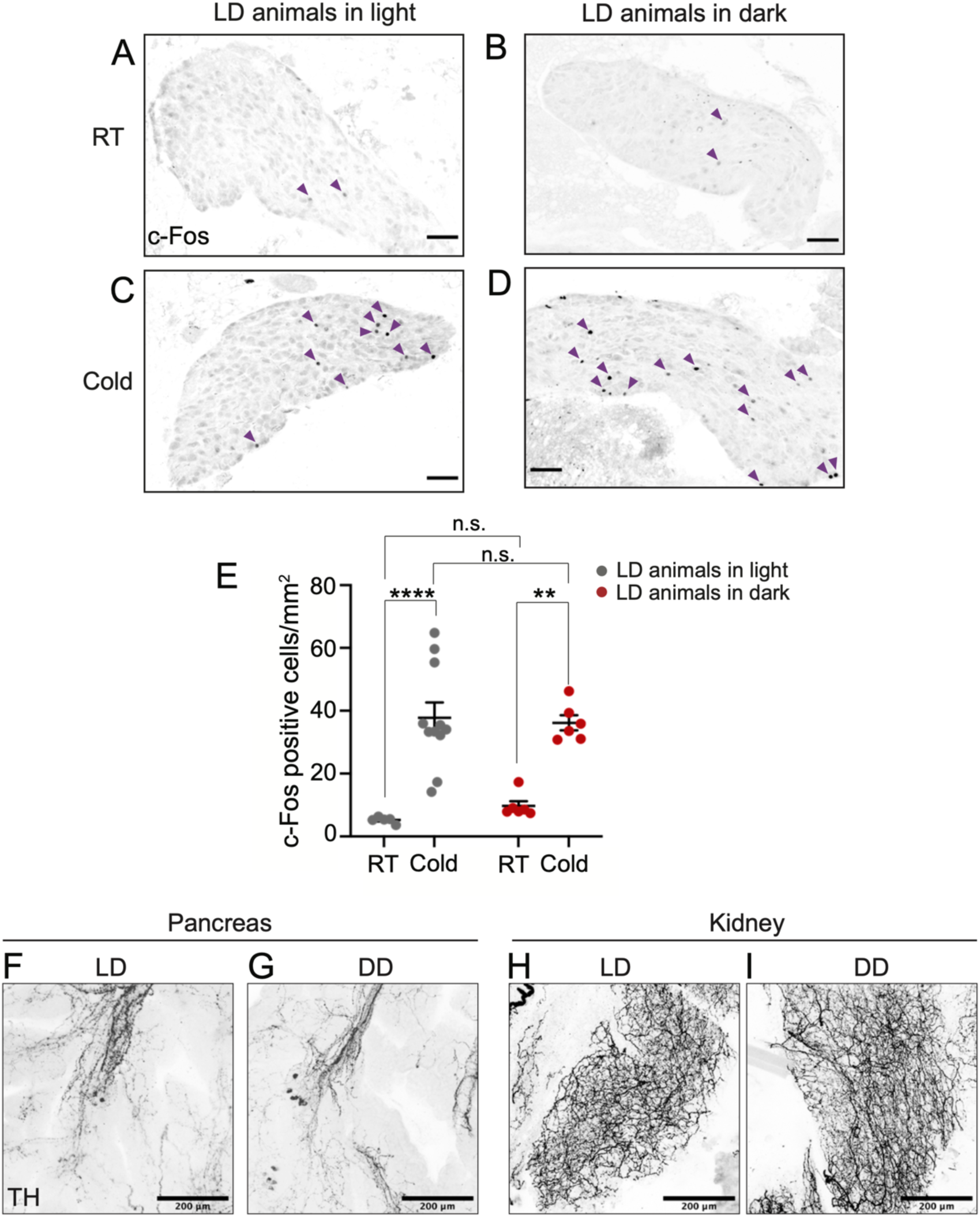
Additional analyses of sympathetic activity and innervation. **(A-D)** c-Fos immunostaining in the celiac-superior mesenteric ganglion complex (CG-SMG) of LD animals that are exposed to a cold stimulus (4°C, 1 hr) and at room temperature (RT) in the light (re-plotted from figure 3 A, C for comparison) or dark. Scale bar: 100μm. **(E)** Quantification of c-Fos-positive sympathetic neurons at RT and in response to cold exposure (4°C, 1 hr) in LD animals kept in the light (re-plotted from figure 3E) or dark during the cold exposure or at RT. Cold exposure significantly increased the number of c-Fos-positive neurons to a similar level for LD animals kept in the light or dark during the cold exposure. Data are presented as means ± s.e.m with n=6 mice for each group. ****p<0.0001, unpaired t-test. **(F-I)** Immunostaining for tyrosine hydroxylase (TH) followed by wholemount clearing shows that sympathetic innervation of peripheral organs, the pancreas **(F, G)** and kidney **(H, I)**, are unaffected by dark rearing. Scale bar: 200μm.

**Fig. S4.**
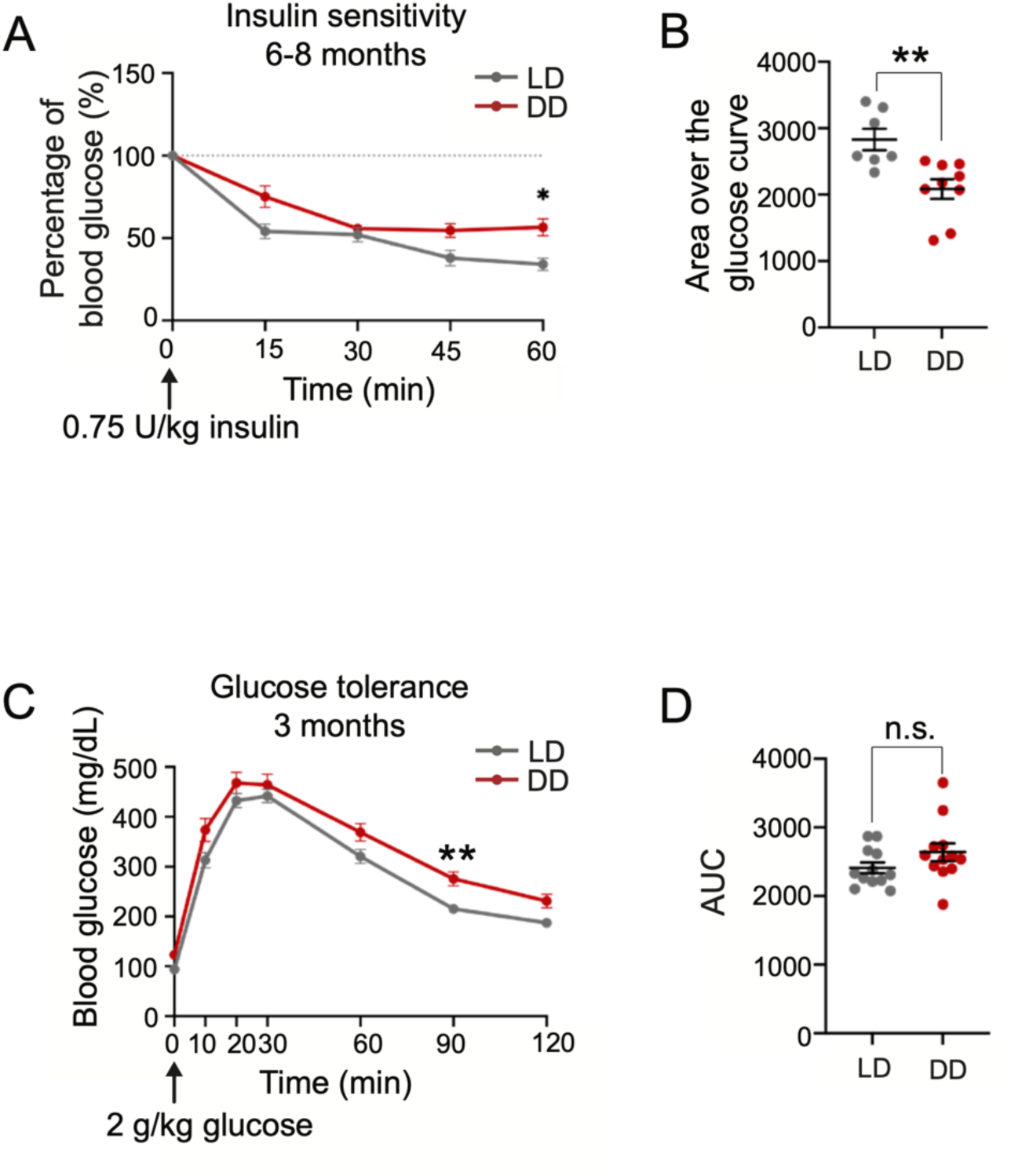
Additional analyses of metabolic functions in older dark-reared mice. **(A,B)** Insulin sensitivity tests. Dark-reared male mice show pronounced insulin resistance at 6-8 months. Data are represented as percentages (%) of the blood glucose level at time “0”. Area over the glucose curve of **(A)** is shown in **(B).** The 100% blood glucose level at time 0 is set as the baseline. Results are means ± s.e.m with n=7 mice for LD and 9 for DD. * p<0.05; **p<0.01, two-way ANOVA, Sidak’s multiple comparisons tests for **(A)** and unpaired t-test for **(B)**. **(C, D)** Glucose tolerance tests. Dark-reared male mice show a mild impairment in glucose tolerance at 3 months. Area under the curve (AUC) of **(C)** is shown in **(D)**. Results are means ± s.e.m with n=12 mice for each group. * p<0.05, two-way ANOVA, Sidak’s multiple comparisons tests for **(C)** and n.s, not significant, unpaired t-test for **(D)**.

## Notes

### Competing Interest Statement

The authors have declared no competing interest.

## References

1. T. A. LeGates, D. C. Fernandez, S. Hattar, Light as a central modulator of circadian rhythms, sleep and affect. Nature reviews. Neuroscience 15, 443–454 (2014).

2. J. Bass, J. S. Takahashi, Circadian integration of metabolism and energetics. Science 330, 1349–1354 (2010).

3. E. Poggiogalle, H. Jamshed, C. M. Peterson, Circadian regulation of glucose, lipid, and energy metabolism in humans. Metabolism: clinical and experimental 84, 11–27 (2018).

4. H. Reinke, G. Asher, Crosstalk between metabolism and circadian clocks. Nature reviews. Molecular cell biology 20, 227–241 (2019).

5. A. Ishihara, A. B. Courville, K. Y. Chen, The Complex Effects of Light on Metabolism in Humans. Nutrients 15, (2023).

6. S. Hattar, H. W. Liao, M. Takao, D. M. Berson, K. W. Yau, Melanopsin-containing retinal ganglion cells: architecture, projections, and intrinsic photosensitivity. Science 295, 1065–1070 (2002).

7. A. D. Guler, C. M. Altimus, J. L. Ecker, S. Hattar, Multiple photoreceptors contribute to nonimage-forming visual functions predominantly through melanopsin-containing retinal ganglion cells. Cold Spring Harbor symposia on quantitative biology 72, 509–515 (2007).

8. L. Lazzerini Ospri, G. Prusky, S. Hattar, Mood, the Circadian System, and Melanopsin Retinal Ganglion Cells. Annual review of neuroscience 40, 539–556 (2017).

9. A. D. Guler et al., Melanopsin cells are the principal conduits for rod-cone input to non-image-forming vision. Nature 453, 102–105 (2008).

10. J. J. Meng et al., Light modulates glucose metabolism by a retina-hypothalamus-brown adipose tissue axis. Cell 186, 398–412 e317 (2023).

11. S. M. Fan et al., External light activates hair follicle stem cells through eyes via an ipRGC-SCN-sympathetic neural pathway. Proceedings of the National Academy of Sciences of the United States of America 115, E6880–E6889 (2018).

12. D. C. Fernandez et al., Light Affects Mood and Learning through Distinct Retina-Brain Pathways. Cell 175, 71–84 e18 (2018).

13. K. Obayashi, K. Saeki, N. Kurumatani, Ambient Light Exposure and Changes in Obesity Parameters: A Longitudinal Study of the HEIJO-KYO Cohort. The Journal of clinical endocrinology and metabolism 101, 3539–3547 (2016).

14. Y. M. Park, A. J. White, C. L. Jackson, C. R. Weinberg, D. P. Sandler, Association of Exposure to Artificial Light at Night While Sleeping With Risk of Obesity in Women. JAMA Intern Med 179, 1061–1071 (2019).

15. K. Nagai, M. Sekitani, K. Otani, H. Nakagawa, Little or no induction of hyperglycemia by 2-deoxy-D-glucose in hereditary blind microphthalmic rats. Life Sci 43, 1575–1582 (1988).

16. M. C. Wanet-Defalque et al., High metabolic activity in the visual cortex of early blind human subjects. Brain research 446, 369–373 (1988).

17. E. Scott-Solomon, E. Boehm, R. Kuruvilla, The sympathetic nervous system in development and disease. Nature reviews. Neuroscience 22, 685–702 (2021).

18. M. G. Myers, Jr., A. H. Affinati, N. Richardson, M. W. Schwartz, Central nervous system regulation of organismal energy and glucose homeostasis. Nat Metab 3, 737–750 (2021).

19. A. Kalsbeek et al., Hormonal control of metabolism by the hypothalamus-autonomic nervous system-liver axis. Frontiers of hormone research 42, 1–28 (2014).

20. B. Ahren, Autonomic regulation of islet hormone secretion--implications for health and disease. Diabetologia 43, 393–410 (2000).

21. K. Nonogaki, New insights into sympathetic regulation of glucose and fat metabolism. Diabetologia 43, 533–549 (2000).

22. E. E. Lin, E. Scott-Solomon, R. Kuruvilla, Peripheral Innervation in the Regulation of Glucose Homeostasis. Trends in neurosciences 44, 189–202 (2021).

23. T. J. Bartness, V. Ryu, Neural control of white, beige and brown adipocytes. Int J Obes Suppl 5, S35–39 (2015).

24. Y. Saito et al., Effect of bright light exposure on muscle sympathetic nerve activity in human. Neuroscience letters 219, 135–137 (1996).

25. A. Niijima, K. Nagai, N. Nagai, H. Nakagawa, Light enhances sympathetic and suppresses vagal outflows and lesions including the suprachiasmatic nucleus eliminate these changes in rats. J Auton Nerv Syst 40, 155–160 (1992).

26. A. Niijima, K. Nagai, N. Nagai, H. Akagawa, Effects of light stimulation on the activity of the autonomic nerves in anesthetized rats. Physiol Behav 54, 555–561 (1993).

27. K. L. Tamashiro, C. E. Terrillion, J. Hyun, J. I. Koenig, T. H. Moran, Prenatal stress or high-fat diet increases susceptibility to diet-induced obesity in rat offspring. Diabetes 58, 1116–1125 (2009).

28. M. M. Glavas et al., Early overnutrition results in early-onset arcuate leptin resistance and increased sensitivity to high-fat diet. Endocrinology 151, 1598–1610 (2010).

29. K. S. Chew et al., A subset of ipRGCs regulates both maturation of the circadian clock and segregation of retinogeniculate projections in mice. eLife 6, (2017).

30. D. Al Rijjal, M. B. Wheeler, A protocol for studying glucose homeostasis and islet function in mice. STAR Protoc 3, 101171 (2022).

31. M. Komatsu, M. Takei, H. Ishii, Y. Sato, Glucose-stimulated insulin secretion: A newer perspective. J Diabetes Investig 4, 511–516 (2013).

32. T. Alquier, V. Poitout, Considerations and guidelines for mouse metabolic phenotyping in diabetes research. Diabetologia 61, 526–538 (2018).

33. S. Virtue, A. Vidal-Puig, GTTs and ITTs in mice: simple tests, complex answers. Nat Metab 3, 883–886 (2021).

34. S. E. La Fleur, Daily rhythms in glucose metabolism: suprachiasmatic nucleus output to peripheral tissue. Journal of neuroendocrinology 15, 315–322 (2003).

35. B. Ahren, G. J. Taborsky, Jr., P. J. Havel, Differential impairment of glucagon responses to hypoglycemia, neuroglycopenia, arginine, and carbachol in alloxan-diabetic mice. Metabolism: clinical and experimental 51, 12–19 (2002).

36. M. Hatting, C. D. J. Tavares, K. Sharabi, A. K. Rines, P. Puigserver, Insulin regulation of gluconeogenesis. Annals of the New York Academy of Sciences 1411, 21–35 (2018).

37. C. C. Hughey, D. H. Wasserman, R. S. Lee-Young, L. Lantier, Approach to assessing determinants of glucose homeostasis in the conscious mouse. Mammalian genome : official journal of the International Mammalian Genome Society 25, 522–538 (2014).

38. F. Karpe, J. R. Dickmann, K. N. Frayn, Fatty acids, obesity, and insulin resistance: time for a reevaluation. Diabetes 60, 2441–2449 (2011).

39. T. J. Bartness, Y. Liu, Y. B. Shrestha, V. Ryu, Neural innervation of white adipose tissue and the control of lipolysis. Front Neuroendocrinol 35, 473–493 (2014).

40. M. Sheng, M. E. Greenberg, The regulation and function of c-fos and other immediate early genes in the nervous system. Neuron 4, 477–485 (1990).

41. R. Candal, V. Reddy, N. S. Samra, in StatPearls. (Treasure Island (FL), 2023).

42. W. Li, G. Yu, Y. Liu, L. Sha, Intrapancreatic Ganglia and Neural Regulation of Pancreatic Endocrine Secretion. Front Neurosci 13, 21 (2019).

43. H. Torres et al., Sympathetic innervation of the mouse kidney and liver arising from prevertebral ganglia. American journal of physiology. Regulatory, integrative and comparative physiology 321, R328–R337 (2021).

44. A. Zsombok, L. D. Desmoulins, A. V. Derbenev, Sympathetic circuits regulating hepatic glucose metabolism: where we stand. Physiological reviews 104, 85–101 (2024).

45. R. Kvetnansky, V. K. Weise, N. B. Thoa, I. J. Kopin, Effects of chronic guanethidine treatment and adrenal medullectomy on plasma levels of catecholamines and corticosterone in forcibly immobilized rats. The Journal of pharmacology and experimental therapeutics 209, 287–291 (1979).

46. Y. Zhang et al., V3 spinal neurons establish a robust and balanced locomotor rhythm during walking. Neuron 60, 84–96 (2008).

47. R. R. Kalyani, J. M. Egan, Diabetes and altered glucose metabolism with aging. Endocrinology and metabolism clinics of North America 42, 333–347 (2013).

48. A. Kalsbeek, S. la Fleur, E. Fliers, Circadian control of glucose metabolism. Molecular metabolism 3, 372–383 (2014).

49. Z. Mirzadeh, C. L. Faber, M. W. Schwartz, Central Nervous System Control of Glucose Homeostasis: A Therapeutic Target for Type 2 Diabetes? Annu Rev Pharmacol Toxicol 62, 55–84 (2022).

50. D. Trico, A. Natali, S. Arslanian, A. Mari, E. Ferrannini, Identification, pathophysiology, and clinical implications of primary insulin hypersecretion in nondiabetic adults and adolescents. JCI Insight 3, (2018).

51. C. Martin, K. S. Desai, G. Steiner, Receptor and postreceptor insulin resistance induced by in vivo hyperinsulinemia. Canadian journal of physiology and pharmacology 61, 802–807 (1983).

52. M. P. Czech, Insulin action and resistance in obesity and type 2 diabetes. Nature medicine 23, 804–814 (2017).

53. D. A. Antonetti, P. S. Silva, A. W. Stitt, Current understanding of the molecular and cellular pathology of diabetic retinopathy. Nature reviews. Endocrinology 17, 195–206 (2021).

54. G. O. de Souza, F. Wasinski, J. Donato, Jr., Characterization of the metabolic differences between male and female C57BL/6 mice. Life Sci 301, 120636 (2022).

55. F. Mauvais-Jarvis, Sex differences in metabolic homeostasis, diabetes, and obesity. Biology of sex differences 6, 14 (2015).

56. F. Mauvais-Jarvis, D. J. Clegg, A. L. Hevener, The role of estrogens in control of energy balance and glucose homeostasis. Endocrine reviews 34, 309–338 (2013).

57. D. M. Berson, F. A. Dunn, M. Takao, Phototransduction by retinal ganglion cells that set the circadian clock. Science 295, 1070–1073 (2002).

58. S. Hattar et al., Central projections of melanopsin-expressing retinal ganglion cells in the mouse. The Journal of comparative neurology 497, 326–349 (2006).

59. J. R. Jones, S. Chaturvedi, D. Granados-Fuentes, E. D. Herzog, Circadian neurons in the paraventricular nucleus entrain and sustain daily rhythms in glucocorticoids. Nature communications 12, 5763 (2021).

60. J. Hu et al., Melanopsin retinal ganglion cells mediate light-promoted brain development. Cell 185, 3124–3137 e3115 (2022).

61. E. Badoer, C. W. Ng, R. De Matteo, Glutamatergic input in the PVN is important in renal nerve response to elevations in osmolality. Am J Physiol Renal Physiol 285, F640–650 (2003).

62. R. M. Buijs, S. J. Chun, A. Niijima, H. J. Romijn, K. Nagai, Parasympathetic and sympathetic control of the pancreas: a role for the suprachiasmatic nucleus and other hypothalamic centers that are involved in the regulation of food intake. The Journal of comparative neurology 431, 405–423 (2001).

63. S. E. la Fleur, A. Kalsbeek, J. Wortel, R. M. Buijs, Polysynaptic neural pathways between the hypothalamus, including the suprachiasmatic nucleus, and the liver. Brain research 871, 50–56 (2000).

64. G. Pail et al., Bright-light therapy in the treatment of mood disorders. Neuropsychobiology 64, 152–162 (2011).

65. M. Sene-Fiorese et al., The potential of phototherapy to reduce body fat, insulin resistance and “metabolic inflexibility” related to obesity in women undergoing weight loss treatment. Lasers Surg Med 47, 634–642 (2015).

66. E. I. Bylesjo, K. Boman, L. Wetterberg, Obesity treated with phototherapy: four case studies. The International journal of eating disorders 20, 443–446 (1996).

67. R. F. Nieuwenhuis, P. F. Spooren, J. J. Tilanus, [Less need for insulin, a surprising effect of phototherapy in insulin-dependent diabetes mellitus]. Tijdschr Psychiatr 51, 693–697 (2009).

68. N. H. Allen et al., Insulin sensitivity after phototherapy for seasonal affective disorder. Lancet 339, 1065–1066 (1992).

69. D. H. Quach, M. Oliveira-Fernandes, K. A. Gruner, W. G. Tourtellotte, A sympathetic neuron autonomous role for Egr3-mediated gene regulation in dendrite morphogenesis and target tissue innervation. The Journal of neuroscience : the official journal of the Society for Neuroscience 33, 4570–4583 (2013).

70. A. M. Ceasrine, E. E. Lin, D. N. Lumelsky, R. Iyer, R. Kuruvilla, Adrb2 controls glucose homeostasis by developmental regulation of pancreatic islet vasculature. eLife 7, (2018).

71. C. B. Wollheim, P. Meda, P. A. Halban, Isolation of pancreatic islets and primary culture of the intact microorgans or of dispersed islet cells. Methods in enzymology 192, 188–223 (1990).

72. H. A. Messal et al., Antigen retrieval and clearing for whole-organ immunofluorescence by FLASH. Nature protocols 16, 239–262 (2021).

